# Multi-reference spectral library yields almost complete coverage of heterogeneous LC-MS/MS data sets

**DOI:** 10.1101/180448

**Authors:** Constantin Ammar, Evi Berchtold, Gergely Csaba, Andreas Schmidt, Axel Imhof, Ralf Zimmer

## Abstract

Spectral libraries play a central role in the analysis of data-independent-acquisition (DIA) proteomics experiments. A main assumption in current spectral library tools is that a single characteristic intensity pattern (CIP) suffices to describe the fragmentation of a peptide in a particular charge state (peptide charge pair). However, we find that this is often not the case. We carry out a systematic evaluation of spectral variability over public repositories and in-house datasets. We show that spectral variability is widespread and partly occurs under fixed experimental conditions. Using clustering of preprocessed spectra, we derive a limited number of Multiple Characteristic Intensity Patterns (MCIPs) for each peptide charge pair, which allow almost complete coverage of our heterogeneous dataset without affecting the false discovery rate. We show that a MCIP library derived from public repositories performs in most cases similar to a “custom-made” spectral library, which has been acquired under identical experimental conditions as the query spectra. We apply the MCIP approach to a DIA data set and observe a significant increase in peptide recognition. We propose the MCIP approach as an easy-to-implement addition to current spectral library search engines and as a new way to utilize the data stored in spectral repositories.

## Introduction

D*ata dependent acquisition* (DDA) approaches are still the standard of proteomics data acquisition. In DDA, selected *precursor ions* are isolated in a small mass window and subsequently submitted for fragmentation and MS measurement^1,2^, giving *MS2 spectra*. The most widely applied DDA approach is also called *shotgun proteomics*, whereby in each duty cycle fragmentation spectra of the N most intense precursor ions are acquired (Top N). The corresponding MS2 data are commonly analyzed by scoring the *mass to charge* (m/z) values of the most intense fragment peaks against a theoretical prediction of m/z values of fragment ions derived from sequence databases^3^. The theoretical m/z values of fragment ions are discriminative as, in most cases, each peak in the MS2 fragmentation spectrum stems from the same precursor ion. Additionally, the m/z value of the submitted precursor ion is known, which narrows down the number of possible matches in the sequence database. Peptide precursors not selected for fragmentation are excluded from the result since sequence confirmation is missing^4^. As precursor ion selection can be described as semi-random^5^, DDA approaches are also problematic for quantification, as a peptide measured in a first sample, might not be identified in a second sample, even though it is abundant.

*Selected reaction monitoring* (SRM, alternatively multiple reaction monitoring (MRM) or parallel reaction monitoring (PRM)) approaches^6,7^ address the problem of reproducibility by a fixed preselection of peptide precursor ions. This approach allows very sensitive and accurate quantitation of a small number of proteins in each LC-MS run, however the overall coverage of the proteome is low due to the preselection. Higher coverage can only be achieved by measuring the sample with multiple precursor lists. Unexpected sample variation, such as post-translational modifications cannot be detected.

*Data independent acquisition* (DIA) approaches try to overcome these limitations by omitting the preselection of precursor ions^8–10^. To reduce spectral complexity, many applications scan MS/MS spectra of medium sized isolation windows (5 – 50m/z) over a wide m/z range^11–17^. In general, the possibilities for spectral searches via sequence databases are challenging for DIA data^18,19^ due to the ambiguity of m/z values in complex peptide mixtures. Thus, many commonly used approaches rely on *spectral libraries,* also considering fragment ion intensities^13,14^. These libraries are obtained from DDA proteomics experiments, by generating a *characteristic intensity pattern* (CIP) of m/z and ion intensity pairs (m,i) from confidently identified MS2 spectra for each peptide in a distinct charge state (peptide charge pair)^20–23^.

A library pattern must be constructed such that it is sufficiently specific (implying few false positives) while maintaining high sensitivity (few false negatives). A library of CIPs is then compared to the measured fragmentation spectrum using a similarity measure. Most current approaches for the construction of library patterns employ the scoring measure *dot product*^24–27^ or the related *spectral contrast angle*^28–30^ as scoring measure. To our knowledge, all tools try to approximate one unique CIP from the available measured fragmentation spectra.

Prior to their use in DIA approaches, spectral libraries have been employed to speed up and increase confidence in peptide recognition^20^ and therefore, large spectral repositories^31–34^ have been compiled.

In current DIA applications like OpenSWATH^14^, *chromatography-based* scores such as retention time are used to find MS2 fragmentation spectra, which are then matched with a library CIP.

Hence, having an accurate spectral library and a highly reproducible and calibrated LC system are key factors determining the quality of a DIA experiment.

In the context of these developments, improved spectral libraries gain renewed importance. In this study, we present a systematic analysis of fragmentation spectra identified with high confidence, by generating and evaluating a model spectral library.

We integrate data from the databases ProteomeTools^36^ (further referred to as Kuster-Set), Pan Human Library^37^ (further referred to as Aebersold-Set) as well as from our own lab (further referred to as Imhof-Set). The Kuster Set contains 8 different combinations of fragmentation type, fragmentation energy and readout, all acquired on an Orbitrap Fusion Lumos mass spectrometer. The Aebersold-Set had fixed fragmentation settings and was acquired on an AB SCIEX TripleTOF 5600+ system from different human tissues and cell lines. The Imhof-Set had fixed fragmentation settings and was acquired on an AB SCIEX TripleTOF 6600 from different organisms and cell lines (see also table 1 for an overview). We only use peptides, which have been measured and identified at least 20 times across several experiments, yielding ≥ 20 *replicate* fragmentation spectra for each peptide charge pair. This gives us a statistical impression of the fragmentation for each peptide.

**Table 1.** Overview over the datasets used in this study and the corresponding experimental parameters.

| id | Intrument | Readout | Fragmentation | Background Matrix | Lab | Collision Energy |
| --- | --- | --- | --- | --- | --- | --- |
| L_HCD_O_20 | Orbitrap Fusion Lumos | Orbitrap | HCD | synthetic peptides | Kuster | 20% |
| L_HCD_O_23 | Orbitrap Fusion Lumos | Orbitrap | HCD | synthetic peptides | Kuster | 23% |
| L_HCD_O_25 | Orbitrap Fusion Lumos | Orbitrap | HCD | synthetic peptides | Kuster | 25% |
| L_HCD_O_28 | Orbitrap Fusion Lumos | Orbitrap | HCD | synthetic peptides | Kuster | 28% |
| L_HCD_O_30 | Orbitrap Fusion Lumos | Orbitrap | HCD | synthetic peptides | Kuster | 30% |
| L_HCD_O_35 | Orbitrap Fusion Lumos | Orbitrap | HCD | synthetic peptides | Kuster | 35% |
| L_HCD_I_28 | Orbitrap Fusion Lumos | Ion Trap | HCD | synthetic peptides | Kuster | 28% |
| L_CID_I_35 | Orbitrap Fusion Lumos | Ion Trap | CID | synthetic peptides | Kuster | 35% |
| Q_CID_AEBERSOLD | AB SCIEX TripleTOF 5600+ | TOF Analyzer | CID | Human tissue + Cell lines | Aebersold | rolling |
| Q_CID_IMHOF | AB SCIEX TripleTOF 6600 | TOF analyzer | CID | Drosophila tissue + Cell lines | Imhof | rolling |

We first demonstrate, that a surprisingly large fraction of MS2 spectra corresponding to the same peptide charge pair is strongly heterogeneous across experimental conditions. This heterogeneity represents a large drawback of using public repositories for spectral library searching, which are mostly obtained under different experimental conditions than the query spectra they are used on. A common practice in many proteomics laboratories is hence the generation of custom-made spectral libraries, especially in the context of DIA experiments^38^. This means it is necessary to generate a spectral library from DDA runs of the desired sample under as similar experimental conditions as possible. One obvious problem is the experimental and computational effort that has to go into creating a custom made library. Additionally, the set of peptides contained in a custom made library is usually orders of magnitude smaller than the peptides available in online repositories.

Based on our findings, we propose the *Multiple Characteristic Intensity Pattern (MCIP)* approach, which is similar to the SpectraST approach by Lam et al.^23^, but differs with respect to the following points: (i) SpectraST uses semi-raw (.mzXML) fragmentation spectra for the generation of spectral libraries, without further preprocessing^23^. We conduct our library generation on MaxQuant^35^ preprocessed peptide identifications without modifications and consider only b- and y-ions (with molecular losses). (ii) As we use preprocessed spectra, we can apply either a ranking prior to the clustering, or use an unranked approach. In both cases, we apply a systematic clustering until all spectra are contained in a cluster and retain all clusters involved. (iii) We determine one CIP from each cluster. This can yield more than one CIP per peptide charge pair.

We compare a spectral library generated with the MCIP approach from a repository with a custom made spectral library and show comparable performance for most datasets. An overview over the major steps taken in this study is given in Fig. 1.

**Figure 1.**
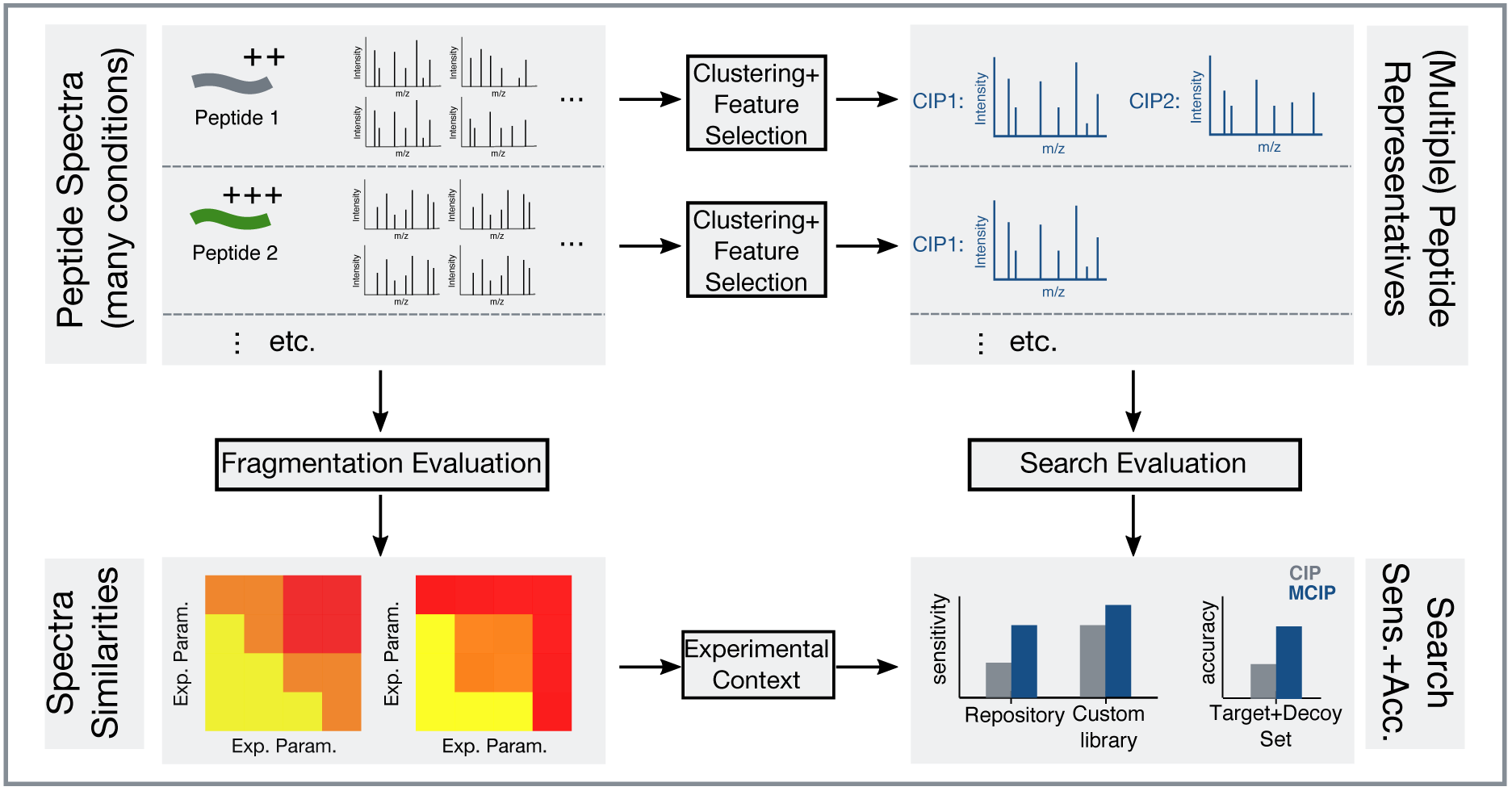
MCIP analysis pipeline. Preprocessed peptide spectra are collected from many datasets and MS runs. The similarities of the spectra are compared over different public repositories and in-house datasets. Multiple Characteristic Intensity Patterns (MCIPs) are generated from the spectra. The search performance (sensitivity, accuracy, etc.) is evaluated in different cross-validation settings, also considering the different experimental contexts.

The MCIP method outperforms the current single CIP approach employed in spectral library searching. We suggest this easy to implement “one-size-fits-all” method as a new way to utilize the data available in spectral archives.

## Experimental Section

### Proteomic analysis via LC-MS/MS on Q-TOF mass spectrometer

Samples were injected into an Ultimate 3000 HPLC system (Thermo Fisher Scientific). For nano-reversed phase separation of tryptic peptide mixtures before MS analysis, peptides were desalted on a trapping column (5 × 0.3 mm inner diameter; packed with C18 PepMap100, 5 μm particle size, 100 Å pore diameter, Thermo-Fisher Scientific). The loading pump flow of 0.1 % formic acid (FA) was set to 25 μl/minute with a washing time of 10 min under isocratic conditions. Samples were separated on an analytical column (150 × 0.075 mm inner diameter; packed with C18RP Reposil-Pur AQ, 2.4 μm particle size, 100 Å pore diameter, Dr. Maisch) using a linear gradient from 4 % to 40 % B in 170 min with a gradient flow of 270 nl/minute. Solvents for sample separation were A 0.1 % FA in water and B: 80 % acetonitrile (ACN), 0.1 % FA in water. The HPLC was directly coupled to the 6600 TOF mass spectrometer using a nano-ESI source (both AB Sciex). A data-dependent method was selected for MS detection and fragmentation of eluting peptides comprising one survey scan for 225 ms from 300 to 1800 m/z and up to 40 tandem MS scans for putative precursors (100-1800 m/z). Precursors were selected according to their intensity. Previously fragmented precursors were excluded from reanalysis for a timespan between 10 and 50 seconds, depending on the experiment (see supplemental table 2). Rolling collision energy setting was enabled, which performs fragmentation at optimized collision energy for the peptide charge pairs. Precursor charge states from +2 to +5 were specifically detected.

### Data analysis of data-dependent LC-MS/MS experiments

The Aebersold-Set and the Imhof Set were analyzed with MaxQuant (version 1.5.1.2 and higher) using the Andromeda search engine^39^ and a sample specific protein database in FASTA format (see supplemental table 1). The settings for database search were as follows: fixed modification carbamidomethylation of cysteine, variable modifications oxidation of methionine and acetylation at the protein N-terminus; ∆mass = 30 ppm for precursors for the first search and 6 ppm for the second search, ∆mass = 60 ppm for TOF fragment ions, enzyme trypsin with specific cleavage and max. two missed cleavages. Peptide hits required a minimum length of 7 amino acids and a minimum score of 10 for unmodified and 40 for modified peptides. Peptide Spectrum Match (PSM) false discovery rate (FDR) was set to 1%.

MaxQuant preprocessing included mass-centroiding of peaks and corresponding intensity adaption, de-isotoping and detection of co-fragmented peptides^35^. The results were returned as msms.txt files, containing the relevant spectral information of fragment ion intensities, retention times, fragment masses as well as charge and modification states of the identified peptide.

The MS proteomics data of the Imhof-Set, including MaxQuant results have been deposited to the ProteomeXchange Consortium (http://proteomecentral.proteomexchange.org) via the PRIDE partner repository with the data set identifiers PXD005060, PXD005063, PXD005100, PXD005111, PXD006245 and PXD006691.

For the Kuster-Set, the MaxQuant files were directly downloaded from the PRIDE repository PXD004732. The raw data for the Aebersold-Set was downloaded from the PRIDE repository PXD000953. More details on the datasets used are displayed in supplemental table 1.

### Library generation settings for the in-house dataset

For the Imhof-Set, the spectral library was generated from DDA data only, with the explicit runs marked in supplemental table 1. For the instrument, the standard configurations as recommended by Sciex were applied to all setups with the vast majority of parameters fixed between all runs. Different settings were only applied to the parameters “Exclude for:” (range 10s - 50s), “Mass tolerance:” (15ppm – 50ppm), “Switch After” (30 spectra – 40 spectra) and “With intensity greater than” (100 - 150). Rolling collision energy was set in all cases. The specific parameters for each input sample are listed in supplemental table 2.

### Selection of processed fragmentation spectra

Peptides were separated by charge into *peptide charge pairs*, because differences in the charge state significantly alter the fragmentation pattern (see supplemental Fig. S1). Only peptide charge pairs, which had at least 20 replicate spectra (see supplemental Fig. S2), were included, to enable the statistical analysis of repeated fragmentation of chemically identical peptides. In our main analysis, we restricted our MCIP approach to only b- and y-ions in charge states up to 2+ with different molecular losses (examples: b3, y4-NH_3_, y6(2+), b5(2+)-H_2_O). Modified peptides were excluded.

### Import of raw fragmentation spectra

To quantify the impact of using all peaks without filtering, an additional analysis with raw spectra was carried out. To assess the influence of the preprocessing method, two different methods of preprocessing the data were applied. In the first approach, the raw spectra were imported from the MaxQuant .apl files contained in the “andromeda” folder in the MaxQuant output folder. We parsed these files and extracted a list of m/z values with corresponding intensities, without b- and y-ion annotation for each spectrum. The spectra were assigned to their respective MaxQuant identification via the spectrum index. In the second approach, the raw .wiff files were processed into the .mzXML format with the MSConvert tool^40^ without any additional filters (yielding profile data), parsed and assigned to the respective MaxQuant identification via the spectrum index. The influence of raw spectral scoring can be seen in supplemental Fig. S3, with an overall lower performance compared to the MaxQuant approach.

### Assessment of the similarity of fragmentation spectra

The similarity among spectra of the same peptide charge pair (*replicate* spectra) can be used as a measure to characterize the fragmentation behavior of peptide charge pairs. As spectra are vectors of (m/z, intensity)-pairs, they can differ in the m/z-values (different peaks) or their intensities, or both.

To assess similarity between replicate spectra, all replicate spectra (at least 20, see previous section) available for a peptide charge pair were compared pairwise to each other.

Each fragmentation spectrum was represented as a *normalized replicate fragmentation vector* (NRFV) *I* = (*i*_1_,*i*_2_, …, *i*_3_), with i_1_ to i_n_ denoting the intensities in the pattern and the indices of the vector implicitly denoting the different fragmentation ions (m/z values). To get vectors of equal length, each fragmentation ion with intensity >0 in any of the replicate spectra was included in every vector. Imputed 0 values were used, if a corresponding ion was not observed. For raw spectra (supplemental Fig. S3) a best bipartite matching method was used. Only vectors with at least four non-zero values (n >4) were used. Each vector was normalized to length |*I*| = 1 (unit vector).

After determining which intensities were included in the NRFVs, the *spectral similarities* between all NRFVs of a peptide charge pair were assessed in a pairwise fashion. For each pair of vectors *X* and *Y* of NRFVs, the *dot score* was calculated using the dot product *similarity measure DP*, defined as

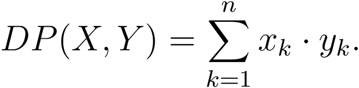

with x_k_ and y_k_ denoting the k-th element of X and Y, respectively.

A pair of fragmentation spectra was called similar, if the dot score of their two corresponding NRFVs was larger than a predefined *similarity threshold* (see below).

### Centroid clustering and CIPs

A central goal of this study is to find a minimal set of *characteristic intensity patterns* (CIPs), able to characterize all observed fragmentation spectra of a peptide charge pair. In order to derive these, a *centroid clustering* approach was employed to determine clusters of similar NRFVs. For each NRFV, the *neighborhood* (all fragmentation spectra with a similarity score greater than the chosen similarity threshold) was determined. The medoid NRFV, corresponding to the spectrum with the best signal to noise ratio (defined via the average intensity of the 2^nd^ to 6^th^ highest peak divided through the median of the remaining peaks) was defined as a CIP, analogous to the SpectraST approach^23^. Additionally, also NRFVs with the largest number of neighbors were defined as CIPs. If not all NRFVs were neighbors to this CIP, it becomes a cluster with all its neighbors and the procedure was repeated on the remaining NRFVs.

Depending on the number of CIPs resulting from this procedure, each peptide charge pair was assigned either a *single CIP* (all spectra of a peptide charge pair assembled in a single cluster) or *multiple CIPs* (MCIPs). The CIPs were referred to by size of their respective cluster: CIP_1_ corresponds to the largest cluster, CIP_i_ to the i-th largest cluster.

### Spectral coverage

The *spectral coverage* was introduced as a measure for the sensitivity of the approach. A spectral library was constructed with the entries for each peptide charge pair consisting either of a single CIP of the largest cluster, or of MCIPs {*CIP*_1_, *CIP*_2_, …, *CIP_n_*} of the n largest clusters.

The single CIP or each element of the MCIPs {*CIP*_1_, *CIP*_2_, …, *CIP_n_*} was then compared to all NRFVs of the peptide charge pair using the dot score. If the dot score was above the similarity threshold for any of the CIPs, the respective spectrum was marked as covered. The spectral coverage denotes the fraction of replicate spectra covered.

### Comparison to custom made spectral libraries

To compare the performance of a custom made library with a MCIP library, we implemented a test set and three training sets. For each experimental setup S, we selected all peptide charge pairs with at least 10 spectra in setup S (and at least 10 spectra in other setups).

Five spectra belonging to S where randomly assigned to the *test* set. The remaining spectra of S were assigned to the first training set, termed the *custom training set*. All spectra that did not belong to S were assigned to the *MCIP training set*. Additionally, the *MCIP custom training set* was defined which consisted of the MCIP training set together with the custom training set. Hence, the custom training set corresponded to the scenario of a custom made spectral library, the MCIP set corresponded to the scenario of having a spectral repository and data measured under differing experimental conditions and the custom MCIP set corresponded to the scenario of integrating a repository library with a MCIP library. Only the main CIP was determined from the custom training set and MCIPs (and also one CIP as a control) were determined from the MCIP training sets. The dot scores of the respective CIPs/MCIPs with the test set were computed.

### Comparison with SpectraST

A comparison of the spectral coverage with the popular SpectraST search engine^23^ was carried out. For this, input files in .pep.XML format suitable for SpecraST were created from the MaxQuant spectrum identifications. Hence for each training set belonging to a specific training-and test set combination, a set of .pep.XML files was generated that contained only the spectra of the specific training set. SpectraST library spectra were then generated from these .pep.XML files. This ensures that the comparison between the MCIP approach and SpectraST approach is carried out with exactly the same underlying data.

To generate the SpectraST library spectra, .pep.XML output files were submitted to SpectraST in library create mode using the default configurations. The resulting raw library was processed to a consensus library using the corresponding SpectraST option. The consensus library mode was chosen, because it has been shown to give the highest number of positive identifications^20^. The consensus library was then quality filtered using the highest quality level (option -cL5) in SpectraST. The raw spectra from the Kuster Set were converted into .mzXML format with the tool MSConvert^40^ and the .mzXML files were subsequently searched with SpectraST.

### Benchmarking via cross validation

To conduct performance testing, a cross validation approach was used. The replicate spectra of each peptide charge pair were split into two fractions. The first fraction consisted of 20% of the spectra and each spectrum was assigned a decoy spectrum 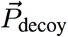 which contained m/z-shuffled intensities of the original spectrum. By shuffling the spectra, the total intensity and the m/z values were preserved, while the spectrum represented changed completely. A 1:1 mixed test set containing original and decoy spectra was then generated. The second fraction consisted of the remaining 80% of spectra. On this fraction, CIP(s) were created as described in the previous sections. The CIP(s) were then similarity scored against the test set using the dot score. A similarity score below the similarity threshold for an original spectrum 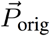 was marked as false negative, a score above the threshold with a decoy spectrum 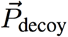 was marked as a false positive. The m/z-shuffling approach is similar to the method employed by Lam et al.^43^, where counting of decoy matches is used library wide to estimate the FDR. Each set of replicate spectra was individually checked via five-fold cross validation in this study. This allowed estimating the relative fractions of false positives and false negatives per peptide charge pair, rather than library wide.

### Choosing a global similarity threshold

A *global similarity threshold* of dot score 0.6 was adapted from the SpectraST search engine^23^ and was subsequently tested using the sampling approach discussed above. This was done to check, whether this threshold would give overall discriminative results.

Each spectrum in the dataset was represented as a NRFV and assigned 1000 differently shuffled decoy vectors. Each NRFV was then dot scored against each decoy vector, which resulted in a distribution of 1000 *shuffled dot scores* for each NRFV. From each distribution of shuffled dot scores, a *local discriminative dot score* was extracted, such that less than 5% of the shuffled dot scores were above this threshold (in other words, the 95% quantile was extracted). Thus, that the use of this dot score would result in 5% acceptance of decoy spectra for a particular NRFV. All locally discriminative dot scores were collected. From the distribution of locally discriminative dot scores, again the 95% quantile was extracted (see supplemental Fig S4). This 95% quantile was 0.62 in this study, which agreed well with the global similarity threshold of 0.6. The approach of extracting two quantiles was taken, because the distribution of shuffled dot scores varied distinctly for different spectra. Hence, taking only one quantile on the distribution of all shuffled dot scores of all spectra combined would result in some spectra (the spectra with generally large shuffled dot scores) being ambiguous. Still, a dot score cutoff of 0.6 might be comparably low considering current high-resolution data.

### Processing of targeted LC-MS/MS runs for CE and isolation window study

Due to the targeted data acquisition setup, the output of the LC-MS/MS experiments was not accessible to standard DDA processing via MaxQuant. The .wiff files were converted to .mzXML using MSConvert (38) and the .mzXML files were then processed using an in-house scoring method, termed *ReScore*. ReScore is based on the scoring described in the publication of the MaxQuant search engine Andromeda^39^. Using DDA runs that were carried out along with the targeted LC-MS/MS runs on the same standardized HeLa Pierce lysate (PXD006691), the scores were compared with Andromeda. The scores show strong correlation with the Andromeda scoring and the vast majority of Andromeda scores is higher than the corresponding ReScore (supplemental Fig. S5). Hence, a certain ReScore cutoff can be used as a reliable cutoff for the Andromeda score.

## Results

### Spectral variability is widespread over experimental conditions

We considered the datasets listed in table 1, containing a total of 10 different experimental settings. We chose a subset of experiments for the Kuster Set with large peptide overlap with either the Aebersold or the Imhof Set, resulting in a heterogeneous set of experimental conditions (see supplemental Fig. S6). To obtain a more detailed understanding of spectral variability, we sorted all replicate spectra corresponding to their respective experimental condition. We then combined all possible pairs of experimental conditions, resulting in 45 pairs (one example pair: Orbitrap Fusion Lumos in HCD mode at CE 25 vs. Sciex Q-ToF 5600+ using CID and optimized rolling collision energy). We assessed the dot scores between the experimental conditions in a pairwise manner. The median values of the resulting dot score distributions are displayed in supplemental Fig. S7 and show a clear clustering after experimental settings. To visualize dissimilar clustering, we plotted the lower 10% quantiles of the pairwise dot score distribution in Fig. 2a.

**Figure 2.**
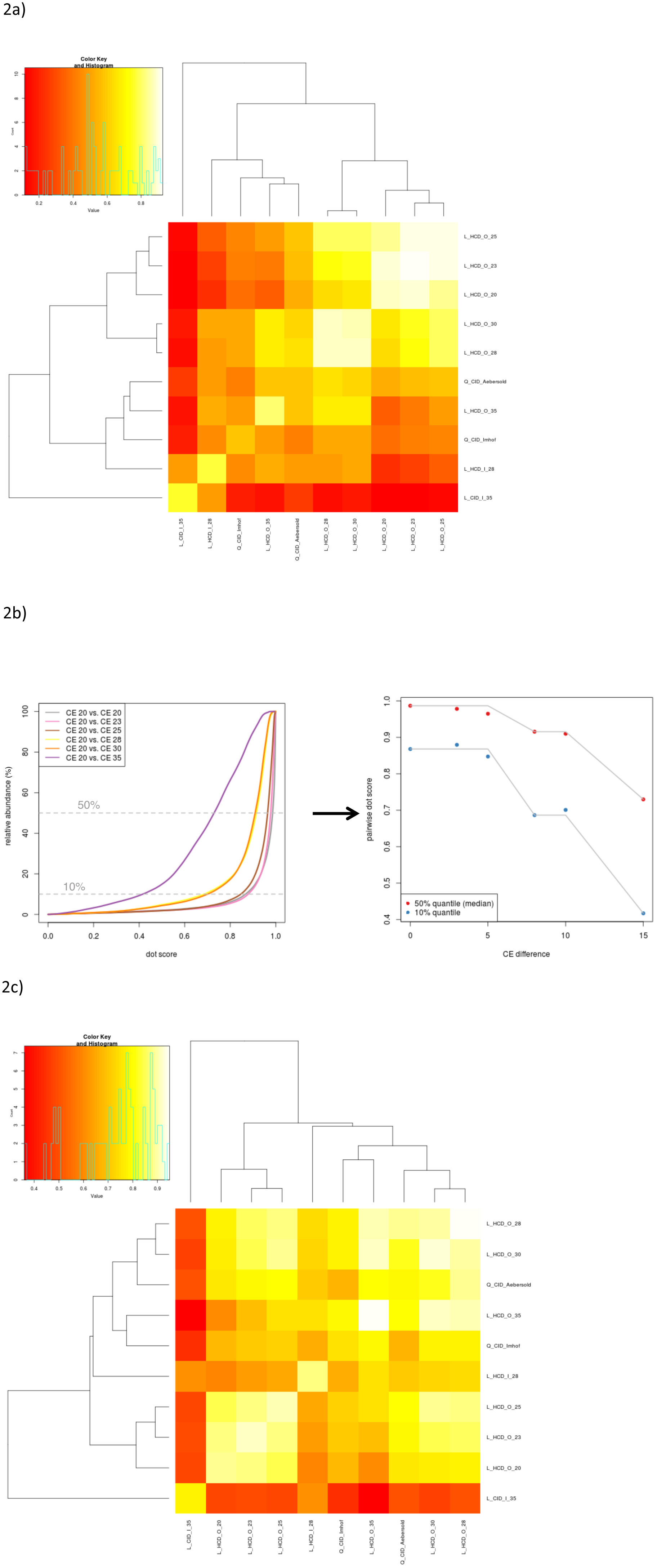
Spectral similarity between experimental conditions. a) Ten percent quantiles of the distributions of dot scores between the different experimental conditions. b) Distribution of pairwise dot scores for the Orbitrap HCD conditions, with the collision energy (CE) 20 setting (left). Median and 10% quantile are drawn in as dashed lines and plotted against the CE difference (right). Two clear drops in similarity are visible. c) Fraction of peptides with no missing spectra after a spectral library search, determined between the different experimental conditions. Depending on the condition pair, the fraction can go down to around 35%.

We observe a large spread in the distributions of dot scores, with visible dependence on the experimental settings. The calculated dot scores are most stable for Orbitrap data generated by HCD fragmentation with collision energies (CEs) from 20 to 30. The Kuster CID@CE35 setup with low-resolution ion trap readout differs most from the remaining setups. The Q-ToF datasets cluster together with the highest CE Orbitrap dataset. We see relatively low dot scores within identical experimental settings (diagonal of the heatmap) for the Q-ToF datasets and for high collision energies as well as for low-resolution readout in the Kuster sets. This underlines, that even under fixed experimental conditions, fragmentation can vary. Some examples are shown in supplemental Fig. S8 and more interpretation of this phenomenon is given in the discussion section.

To investigate the influence of CE on spectral similarity, we considered the distributions of pairwise dot scores between the Orbitrap set at CE 20 and the Orbitrap sets at higher CE (Fig. 2b, left). We see a clear influence of the CE difference on the pairwise distributions. We extracted the 10% quantile and the median from these distributions and plotted them against the CE difference (Fig. 2b, right). We see no visible influence of small CE changes. For CE changes between 5 and 8 we see a clear drop in similarity and between 10 and 15 an even stronger drop, which indicates complex processes underlying peptide fragmentation. Ion trap readout generally shifts the dot scores towards lower values (see supplemental Fig. S9).

To elucidate the drastic effects of the observed variability on peptide identification, we carried out a clustering on the 45 combinations of experimental conditions (Fig 2c). Each combination in the heatmap shows the fraction of peptides that have only one cluster. This corresponds to no spectra clustering outside the main cluster and hence no spectra being missed in a spectral library search. We see that - depending on the experimental combination – as little as 35% of peptides fulfill this condition, with most combinations ranging from 60% to 90%. From these observations, we conclude two main points: (i) The machine setup can play a crucial role for spectral recognitions. (ii) Even fixed experimental settings can lead to spectra being missed.

### Usage of MCIPs yields almost complete spectral coverage

To achieve a global impression of the performance, we assessed the spectral coverage (see method section) for all peptide charge pairs. The spectra were either compared with one CIP (in accordance with the current library approaches) or with MCIPs. Fig. 3a shows a striking improvement upon successive integration of more and more CIPs (red up to orange line) until near-complete coverage is reached. The largest gain is visible upon integration of the second CIP (blue line), which corresponds to the second largest cluster. This improvement clearly shows that significantly higher peptide recognition is possible by simply including two representative spectra for a peptide charge pair instead of just one.

### MCIP library performs comparable to custom-made library and enhances custom-library performance

Custom-made spectral libraries can be seen as the gold standard for creating a high performing spectral library^44^. These libraries, however, come with drawbacks compared to spectral repositories, mainly due to the effort in creating the library or the limited number of peptides in the library.

To evaluate the performance of our libraries, we generated a set of test spectra for each experimental condition. For each set of test spectra, we generated spectral libraries from three different sets of training spectra: 1) The *custom set* only contained spectra measured under the same experimental condition as the test set. 2) The *MCIP* set contained spectra measured under all available conditions *except* the condition of the test set. 3) The *MCIP custom* set contained spectra measured under all available conditions. The MCIP set was chosen in this way to re-create the scenario of having a repository library (the MCIP set) and own data measured under a different experimental setting (the test set).

We then determined either a single CIP or MCIPs from the different sets of training spectra and compared them with the test spectra (see also methods section). This allowed us to directly assess the effect of extending the current singe CIP approach to an MCIP approach.

We first examined the fraction of *missed spectra,* which denotes spectra that would not be detected in a spectral library search, when using a (rather low) similarity threshold of 0.6 (Fig 3b). Using MCIPs always performs better than the current single CIP approach. As a general trend, we see that differently clustering spectra are more common in experimental setups where either the spectral resolution is low (ion trap), or fragmentation energy settings are high. The most challenging setup Kuster CID@CE35 with ion trap readout, leads to around one third of spectra being missed when using a single CIP approach. Integrating multiple CIPs reduces the missed fraction by a factor of 2. Using a custom library further reduces the missed rate to around 5%. Using the MCIP custom training set yields an overall missed rate of 3%. For the other experimental conditions, the MCIP approach gives a similar performance as the custom library approach. In around half of the cases the custom library approach is slightly better, in the other half the MCIP approach performs slightly better. The MCIP approach in combination with the custom approach always increases accuracy. The results described also hold for maximum neighbor clustering (see supplemental Fig. S10) and are stable for different training sets.

An example of a spectrum being detected by a CIP outside the main cluster is given in Fig. 3c. To give an intuitive visualization of the similarities between the spectra, the fragment ion intensities are connected via lines. We see that CIP1, which was acquired at HCD@CE23 has a significantly less prominent b6 ion, which significantly alters the shape of the fragmentation profiles for the higher energy CID@CE35 and HCD@CE28 spectra. The annotated raw spectra corresponding to Fig. 3c are displayed in supplemental Fig. S11.

### Direct Comparison with SpectraST shows significantly increased sensitivity

To further assess the performance of our MCIP approach, we carried out a comparison with the SpectraST^23^ spectral search engine, which is among the most popular in the field^45^. We again determined the fraction of missed spectra at a similarity threshold of 0.6. We generated the SpectraST library on the identical spectra as our own library (see also methods section). We see that the MCIP approach outperforms the single CIP approach of SpectraST in terms of sensitivity in the custom setup as well as in the non-custom setup (Fig 3d).

**Figure 3.**
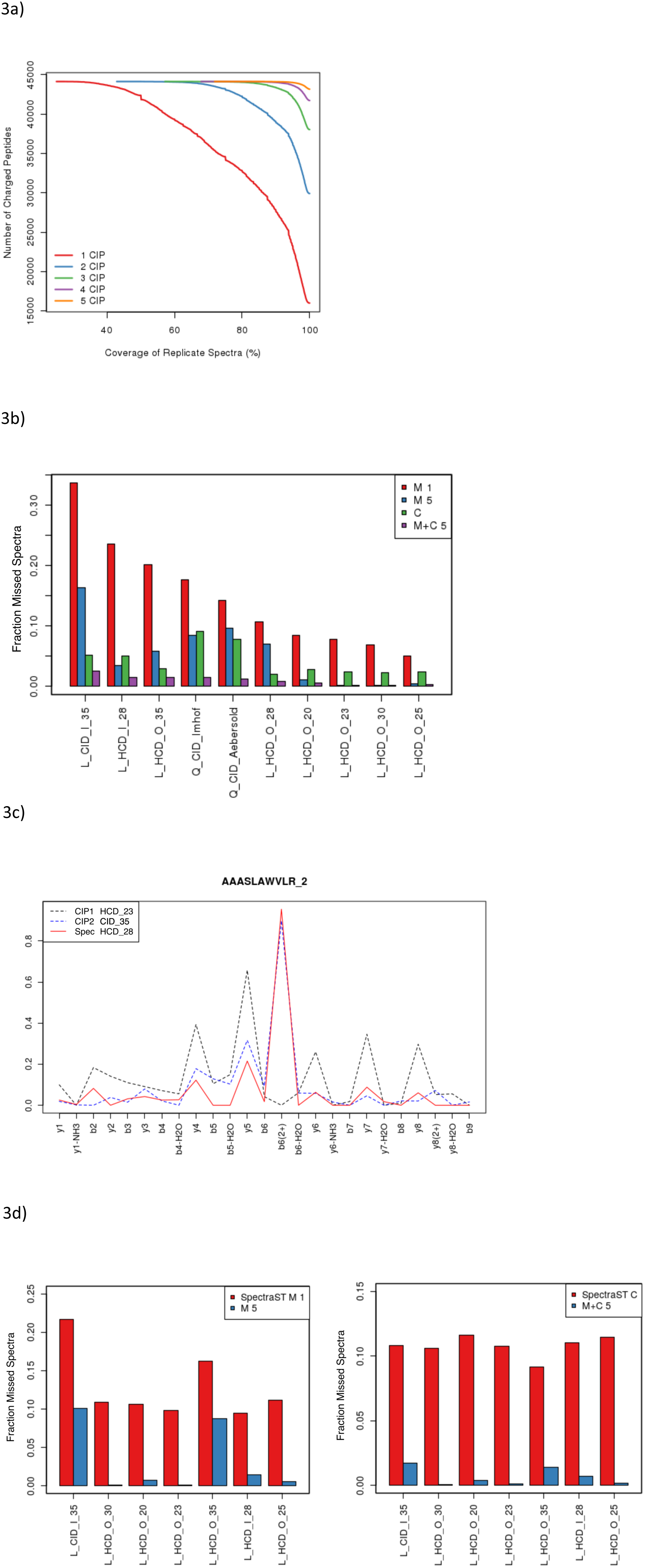
Comparison of the MCIP approach with current approaches. a) Spectral coverage of the whole dataset for different numbers of characteristic intensity patterns (CIPs) integrated in the spectral library. The number of peptide charge pairs (y-axis) is displayed, for which the spectral coverage is larger or equal to the value denoted on the x-axis. The single CIP approach (red line) leaves a large fraction of spectra uncovered. Integrating one more CIP (blue line) into the library gives a strong increase in coverage with successively smaller increases upon integration of more CIPs until almost complete coverage is reached for up to 5 CIPs per peptide charge pair. b) Fraction of spectra with dot score <0.6 to the CIP/MCIPs (missed spectra). Each group of histograms displays one experimental condition the method is tested on. The different clustering approaches are indicated in the legend: M1 equals the MCIP approach with a single CIP (state-of-the-art approach). M5 equals the MCIP approach with a maximum of 5 CIPs included. C equals the custom library approach and M+C 5 equals the MCIP approach with custom spectra included. c) Example of a query spectrum dissimilar to CIP1 (dot score 0.28) but similar to CIP2 (dot score 0.96). The raw spectra are displayed in supplemental Fig. S11. d) Comparing the MCIP approach with SpectraST using identical training and test sets for the MCIP approach and for SpectraST. The number of spectra missed is significantly lower for the MCIP approach both for the non-custom (left) and the custom (right) approach.

### MCIPs increase sensitivity without affecting specificity

As has been shown in the previous sections, MCIPs are able to cover all replicate spectra for many peptide charge pairs, and, thereby improve sensitivity in spectral searches. However, this might come at the cost of reduced specificity (i.e. increase in false positives). Here we investigate, whether using MCIPs affects the number of false positives and the overall accuracy. We tested this by first generating CIPs on 80% of the replicate spectra and then scoring these CIPs against a mixture of the remaining replicate spectra and shuffled decoy spectra. This allowed distinguishing true positives (match of CIP with replicate spectrum) from false positives (match of CIP with decoy spectrum). The procedure is described in more detail in the methods section. Fig. 4a) shows that the overall accuracy for MCIPs (blue line) increases significantly in comparison to a single CIP (red line). For >99% of peptide charge pairs, the minimum accuracy increases by around 10% when integrating all CIPs available for each peptide charge pair in the spectral library (blue line). In Fig. 5b), we see (as expected from the spectral coverage results) a strong decrease in false negatives upon integration of MCIPs. At the same time, we see that the false positive rate displayed in Fig. 5c) is very mildly affected. The results of integrating all MCIPs instead of 5 MCIPs are virtually identical.

**Figure 4.**
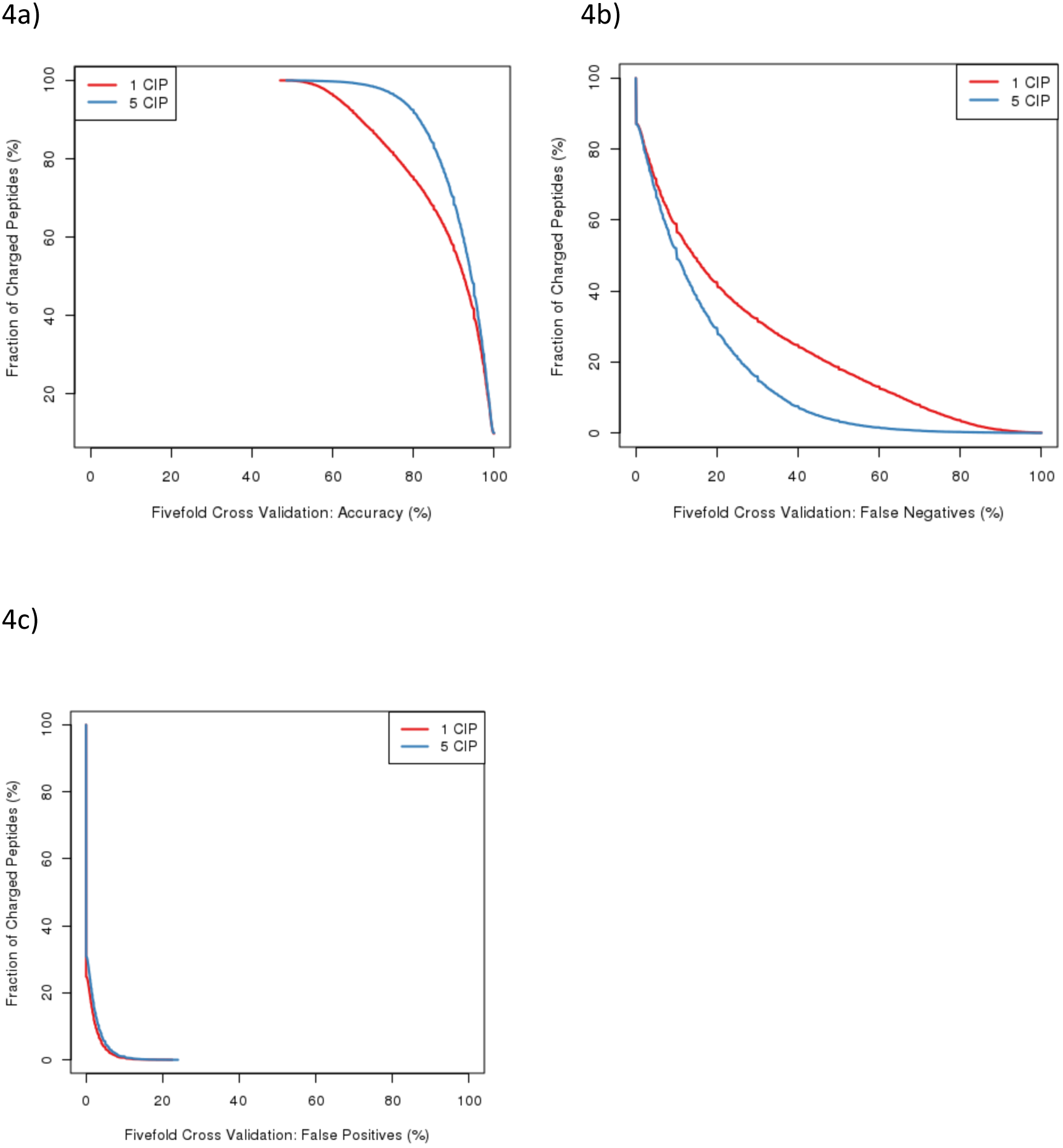
Assessment of accuracy using permuted decoy spectra, as described in the methods section, cumulative plot. a) Comparison of the overall identification accuracy between the single characteristic intensity pattern (CIP) approach (red line) and multiple CIPs (MCIPs) (blue line). A significant improvement upon integration of MCIPs is visible. b) Effect of MCIP integration on false negatives: The false negatives rate is strongly decreased as now also the differently fragmenting ions are integrated. c) Effect of MCIP integration on the false positives rate, which is only marginally increased upon integration of MCIPs.

**Figure 5.**
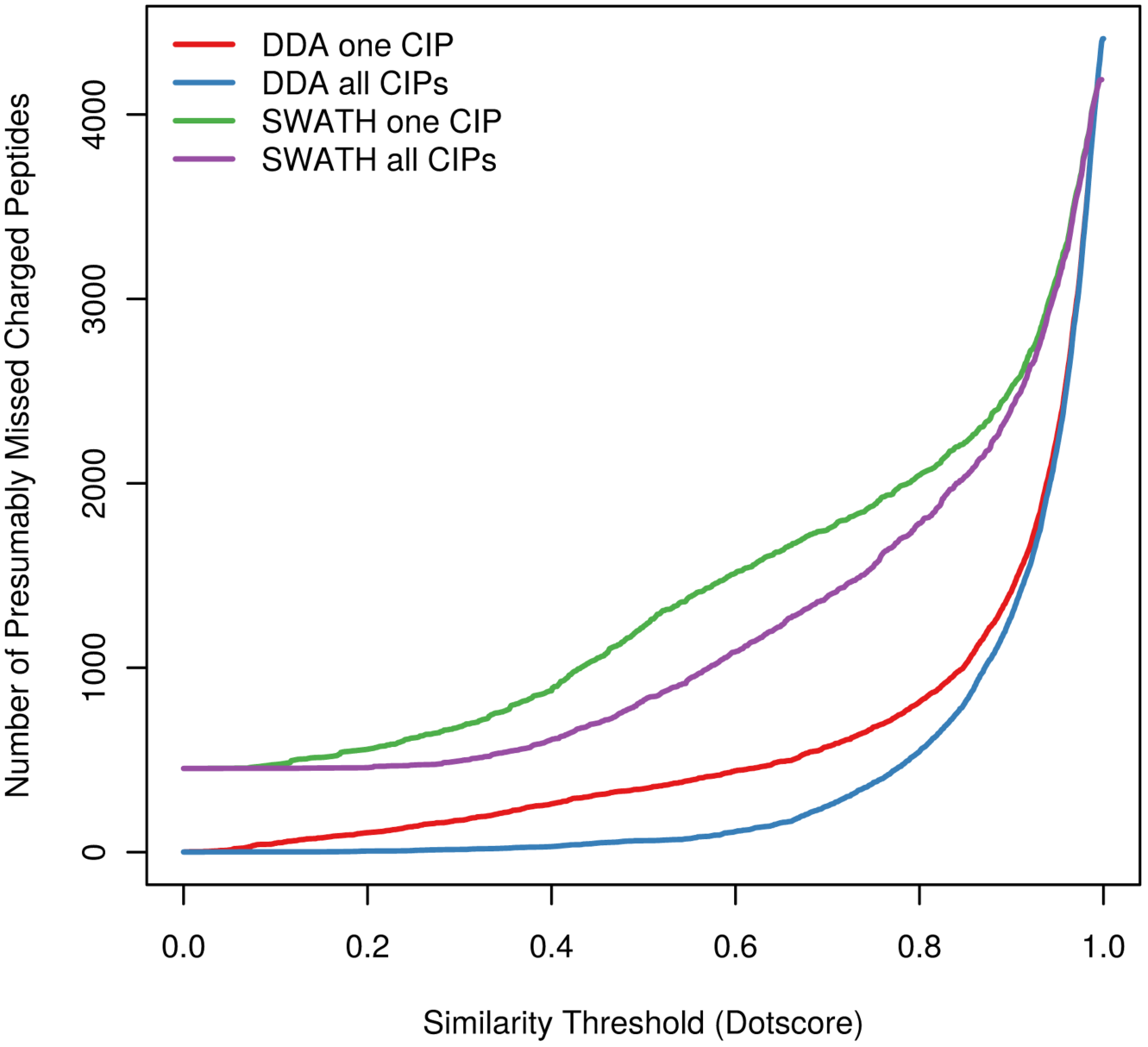
Application of the MCIP approach on SWATH data and comparison with DDA data. For reasonable similarity thresholds, a significant decrease in unidentified peptides can be seen on SWATH data when integrating MCIPs (violet line) in comparison to a single CIP (green line). An analogous behavior is seen for the DDA approach, with a significantly smaller number of missed peptides in both cases (blue line MCIPs, red line single CIP).

### Implications for SWATH data

One of the current applications of spectral libraries is the analysis of SWATH data^14^. In this setup, CIPs are matched with more complex MS2 spectra. We assessed, whether MCIPs can improve the identification rate as compared to using only one CIP.

As described in the methods section, we used a dataset, where the same sample had been identified using DDA and SWATH. We then utilized this setup to derive a spectral library of peptides, which was expected to be in the SWATH data set.

We then searched the library patterns against the DDA run as well as the SWATH run, with the fraction of non-identified spectra (“errors”) plotted against the similarity threshold (Fig. 5). For the DDA run, the results are analogous to the results already presented, with a significant improvement of identification upon integration of MCIPs (red and blue line, respectively).

For the SWATH run, we observe lower baseline identification, with approximately 20% of the patterns not being identified at all, likely due to the higher noise in the SWATH patterns. Nevertheless, also for the SWATH data set, we observe very similar effects when comparing the single CIP approach (green) with the MCIP approach (magenta), with an ^~^30% increase in identification accuracy at reasonable similarity scores (e.g. 29% for a dot score of 0.6). The higher senstivity correspond well with the recent findings that SWATH data analysis is improved when local libraries are used in addition to public libraries^46^.

## Discussion

In our study we have introduced a simple and efficient strategy to deal with heterogeneity of peptide fragmentation. We see that instrument settings can have a huge influence on peptide fragmentation behavior, especially for high-energy and low-resolution spectra. We have shown that exclusion of dissimilar peptide spectra is over-cautious and results in the negligence of many potential hits.

We observe that even under fixed experimental conditions, spectra can vary from each other. This effect is strongly enhanced by low-resolution readout. Additionally, very high collision energy changes also have an effect on differing peptide fragmentation. Unfortunately, it is beyond the scope of our study to fully explain the differences in peptide fragmentation under fixed conditions. However, we carried out some initial screens, using targeted LC-MS/MS runs, where we varied the applied collision energy within the same run. This was done to test, whether a wrong charge state assignment from the machine could account for the effects. Our results show that dot scores are robust over a range of −3V to +3V in most of the cases, which covers differences in collision energy settings caused by wrong precursor charge state assignment (supplemental Fig. S12). We complemented these runs with experiments on the same lysate, where we tried to investigate the influence of the *background matrix* (coeluting peptides/ions in the same isolation window). For this purpose, the precursor isolation width was varied between 1Da and 5 Da. For broader isolation windows, we observed a systematic enrichment in differently fragmenting spectra (supplemental Fig. S12) and an increase in spectral dissimilarity within the same experimental run (supplemental Fig. S13).

For low abundant peptides, it has already been shown that ion interferences with the background matrix can alter the fragmentation spectrum^47^, which we also see in corresponding analyses in supplemental Figs. S13 and S14. Recent studies also show this effect with SWATH-MS data^48^. As a low resolution readout should also strongly increase the effect of interferences, we speculate that the background matrix might be responsible for the observed differences in the fragment spectra.

Our reductionist approach of relying on MaxQuant preprocessed spectra comes at the cost of possibly neglecting important spectral information. The dot score values determined from this approach will be different to the dot score values derived from raw spectra, as the representative vectors are shorter and vector length influences the outcome. Nevertheless, heuristic measures to shorten the vector are applied in common library generation tools^20,22^ and have shown to only mildly affect the overall sensitivity. Additionally, we explicitly tested for accuracy, which is displayed in Fig. 4 of this study.

It should be noted that MaxQuant spectra do not carry fragment ion annotations for fragment ions in charge states larger than 2. To check, whether this affected the outcome of the scoring, we repeated our measurements only on peptides in charge state 2 (which should hence not produce fragment ions with charge larger than 2), with no qualitative differences in the outcome.

Based on our findings, we conclude that even though considerable efforts are being undertaken to extend the amount of available experimental setups in spectral repositories (as for example in the scope of the ProteomeTools^36^ project), this might only be part of the solution. Due to the large variety of machine-setups available, a public library is unlikely to be a perfect fit for the desired setup (including instrument model, fragmentation mode, collision energy settings, fragment ion readout). Additionally, when a well fitting spectral library is found, the user has to constrain to the parameters of the library, which comes at the cost of flexibility in tuning the machine setup. However, even when this is fulfilled, the user is not able to utilize the full amount of spectral data available online, as only the peptides available in the specified setup can be used. As we have shown, using MCIPs frees the user of these constraints and hence improves the usability of spectral resources.

With the advent of quantitative DIA methods like SWATH, the phenomenon of MCIPs becomes important in the context of quantification. If MCIPs are not taken into account, in a significant fraction of cases, fold changes might be miscalculated because peptides that are actually there will be missed because the fragmentation spectrum is different. The increase in spectral recognition of our SWATH data set upon integration of MCIPs (Fig. 5) is a first indication that SWATH benefits from our approach.

## Abbreviations

DIA: data independent acquisition
DDA: data dependent acquisition
SRM: selected reaction monitoring
MRM: multiple reaction monitoring
PRM: parallel reaction monitoring
SWATH: sequential window acquisition of all theoretical mass spectra
CIP: characteristic intensity pattern
MCIP: multiple characteristic intensity patterns
FA: formic acid
TFA: trifluoroacetic acid
ACN: aceonitrile
PSM: peptide spectrum match
cps: counts per second
NRFV: normalized replicate fragmentation vector

